# A functional comparison of readthrough agent ELX-02 across a wide range of nonsense CFTR variants

**DOI:** 10.64898/2026.07.31.740556

**Authors:** Marlies Destoop, Anabela S. Ramalho, Iris A. L. Silva, Annelotte M. Vonk, Sylvia Suen, Marlou C. Bierlaagh, Sylvia F. Boj, Robert G. J. Vries, Jeffrey M. Beekman, Kris De Boeck, Cornelis K. van der Ent, Margarida D. Amaral, Francois Vermeulen, Sacha Spelier, HIT-CF Organoid Study

**Author notes:** S2AQUA—Collaborative Laboratory, Association for a Sustainable and Smart Aquaculture, Avenida ParqueNatural da Ria Formosa s/n, 8700-194 Olhão, Portugal (Iris A. L. Silva).

## Abstract

**Background:** Nonsense variants in *CFTR* account for ~10% of cystic fibrosis (CF) variants and cannot be treated with approved CFTR modulators. Translational readthrough agents such as ELX-02 offer a potential therapeutic strategy, but clinical trials evaluated mainly in G542X CFTR nonsense variant and underlined limited efficacy. This study aimed to evaluate ELX-02-mediated CFTR rescue across a broad range of nonsense variants using patient-derived intestinal organoids (PDIOs) to define variant-specific determinants of readthrough efficacy and assess its potential across a genetically diverse CF population.

**Method:** The ex vivo response to ELX-02 was assessed in 206 PDIOs carrying heterogeneous nonsense variants. CFTR function was quantified using forskolin-induced swelling (FIS) assay after 48-hour exposure to ELX-02. Responses were analysed by genotype and stop codon identity, with secondary validation performed in a selected subset of PDIOs (n = 60).

**Results:** ELX-02–mediated CFTR rescue varied markedly, ranging from responses approaching those observed with approved CFTR modulators (LUM/IVA) to responses at or below detection limit. Overall, maximal responses were modest and at the lower end of the functional range for CFTR modulators. Rescue was dose-dependent and higher in PDIOs carrying two nonsense variants when compared with PDIOs carrying a single nonsense variant combined with a residual or minimal function variant. Nonsense variants in nucleotide-binding domain 1, including G542X, S466X, G550X and R553X, showed relatively higher responsiveness.

**Conclusion:** ELX-02 induces limited and highly heterogeneous CFTR rescue across nonsense variants. PDIO-based functional screening provides a framework to guide patient selection and stratification for future readthrough therapy trials.

## Introduction

Cystic fibrosis (CF) is a life-limiting autosomal recessive disorder caused by variants in the *CFTR* gene, resulting in defective chloride and bicarbonate transport and progressive multiorgan disease, particularly affecting the lungs and gastrointestinal tract (1). Among the more than 2000 described *CFTR* variants, nonsense variants introduce premature termination codons (PTCs), which lead to truncated, non-functional proteins and often trigger degradation of the corresponding mRNAs via nonsense-mediated mRNA decay (NMD) (2-4). Nonsense variants account for 8.4% of all CF-causing *CFTR* alleles worldwide (http://www.genet.sickkids.on.ca/). The subset of people with CF (pwCF) homozygous for a nonsense variant or in trans with another class I variant are ineligible for currently approved CFTR modulators, which primarily target gating or processing defects of the protein.

One promising therapeutic approach for this underserved population involves translational readthrough agents (5). These compounds act by modulating ribosomal fidelity at the PTC site, enabling the incorporation of amino acids (AA) at the site of the PTC and allowing translation to continue to the normal termination codon (NTC), ultimately producing full-length CFTR protein. Mechanistically, different classes of readthrough agents can act through distinct pathways. Aminoglycosides, for example, transiently slow ribosomal translocation and promote mispairing at the nonsense site, whereas more recent compounds, such as di-2,6-aminopurine (DAP), target tRNA-modifying enzymes to facilitate the selective incorporation of cognate AA at the nonsense site (6-9). In all cases, readthrough efficacy is influenced by the identity of the stop codon, the surrounding nucleotide context, and the location of the nonsense within the transcript (10).

Despite their promise, no readthrough agent has yet demonstrated clinical efficacy for pwCF. Ataluren, the furthest developed for Duchenne muscular dystrophy, failed to show robust improvement in large scale CF clinical trials (11, 12). Similarly, ELX-02, a chemically engineered aminoglycoside that reached the furthest stage in CF-specific clinical development, has so far only been tested in a small cohort of patients carrying the G542X variant, either alone or in combination with the CFTR potentiator ivacaftor. Phase 2 trials revealed limited functional and clinical benefit, which may at least partially be explained by suboptimal pharmacodynamic properties, including low pulmonary drug concentrations (6, 13). These translational gaps underscore the challenges of developing readthrough therapies and highlight the need for a deeper mechanistic understanding and broader preclinical evaluation.

Patient-derived intestinal organoids (PDIOs) represent a powerful platform to bridge this translational gap. PDIOs are three-dimensional cultures derived from adult stem cells isolated from intestinal crypts, which recapitulate key features of the native epithelium, including cell polarity, cell-type composition, and CFTR expression predominantly at the apical membrane (14-16). Forskolin-induced swelling (FIS) of PDIOs provides a quantitative readout of CFTR-dependent fluid secretion, which correlates with clinical disease severity, progression, and response to modulators (17, 18). As a result, PDIOs enable patient-specific and variant-specific preclinical assessment of therapeutic strategies, including readthrough compounds, under physiologically relevant conditions (19, 20). Moreover, PDIOs can be expanded and cryopreserved to create living biobanks that represent a wide spectrum of *CFTR* genotypes, facilitating systematic evaluation of novel therapies (21, 22).

Here, we used PDIOs to systematically evaluate the ex vivo response to ELX-02 across a broad set of *CFTR* nonsense variants. By expanding beyond the single G542X genotype previously tested in clinical trials, we aimed to define variant-specific determinants of readthrough efficacy and assess the potential of ELX-02 to rescue CFTR function across a genetically diverse CF population. This study provides a comprehensive preclinical framework to guide future development of readthrough therapies and improve patient stratification for clinical trials.

## Materials and Methods

### PDIO Culture and Expansion

Within the HIT-CF project, 489 PDIOs from PwCF in 12 countries were collected in a central biobank (23) for the HIT-CF Organoid Study (NTR7520). All participating clinical sites were members of the clinical trial network of the European Cystic Fibrosis Society (ECFS-CTN). This study and the accompanying informed consent form were approved by independent ethics committees at each participating site, with final approval for use of all PDIOs by TcBio (24-263). Written informed consent was obtained from each participant and/or the participant’s legal guardian.

PDIOs from patients carrying at least one CFTR nonsense variant were selected for the present analysis. The selected samples were divided between the three European labs participating in the HIT-CF European project (KULeuven, ULisboa and UMC Utrecht). PDIOs were thawed and seeded in a 24-well plate with medium supplemented with Rho-associated kinase inhibitor (RhoKI) according to the previously published Standard Operating Procedure (SOP) (15). Cultures were maintained at 37°C with 5% CO_2_ and passaged weekly to promote growth and budding. PDIO quality and morphology were monitored, and after at least 3 weeks of expansion, cultures were used to perform the FIS assay.

### Forskolin induced swelling assay

PDIO cultures were split mechanically every 7 days and after disruption replated at a density of 20-50 PDIOs per 4 µL droplet in a 96-well plate, as previously described (15). Plates were incubated at 37°C with 5% CO_2_ for a maximum of 10 minutes before adding the respective compound solutions. For each PDIO sample, two screens were performed, with each experiment repeated in duplicate with two technical replicates. In the primary screen, media-only controls were compared to 80 µM ELX-02 and 160 µM ELX-02. In the secondary screen, media-only controls were compared to 40 µM ELX-02, and 80 µM ELX-02.

After a 48-hour incubation period with or without ELX-02, PDIOs were stained with calcein AM for 15-20 minutes to visually assess viability. Following the viability assessment, PDIOs swelling was measured using the FIS assay. PDIOs swelling was stimulated by adding forskolin at a single final concentration of 0.8 µM (no titration). Images were captured every 20 minutes for 120 minutes on a confocal microscope, yielding seven timepoints for analysis. Total organoid area was calculated from the captured images by integrated imaging analysis software (Zen from Zeiss). The raw area data for each timepoint and well were used to calculate the Area Under the Curve (AUC) for normalized swelling over the 120-minute experiment duration. To normalize for differences associated with sites, we characterized ELX-02 mediated swelling of the same G542X/G542X PDIO line at the 3 sites; calculated the mean and normalized all FIS data for the differences of this median at ELX-02 at a concentration of 80 µM. The mean ELX-02 mediated swelling of the G542X/G542X lines were used as a reference line throughout the figures.

### Final dataset and study design

From the HIT-CF biobank, all PDIOs carrying at least one nonsense variant (n = 223) were included in the initial selection for the primary screen. A total of 17 samples were excluded afterwards for the following reasons: reference samples (G542X/G542X) (n = 6), ≥ 1 unknown variant (n = 6), equivocal nonsense variants (G587X/G587X, R347X/R347X) (n = 2), less than 2 CF causing variants (M470V/Q1313X, M470V/W1282X) (n = 2), no nonsense variant (n = 1). After these exclusions, results from 206 PDIOs were available and included in the primary screen.

Based on the results in the primary screen, the highest responders were selected for a subsequent validation screen (secondary screen). In total, 60 PDIOs were re-tested by HUB Organoids: 57 PDIOs identified as top responders and 3 classified as non-responders for negative control. Selection was stratified across three batches, corresponding to the three participating screening laboratories. Sample selection for validation screen was performed by a statistician at UMCU, who was blinded to genotype. Three additional criteria were applied:1) exclusion of PDIOs allocated to the CHOICES clinical trial (part of the HIT-CF project) evaluating diponecaftor (ClinicalTrials.gov NCT064685270; EudraCT 2022-500410-26-01); 2) exclusion of PDIOs with high residual CFTR function (>2500 AUC after forskolin only), as these display limited additional drug-induced swelling; 3) inclusion of the highest 25% of ELX-02 responders at 80 µM (residual function-corrected) at each site.

### Statistics

To evaluate the functional response of each PDIO line to the tested compound solutions, area under the curve (AUC) values were calculated using “R” software. To correct for residual function (swelling induced by forskolin alone indicating partly functional CFTR), the medium-only control was used in which swelling was measured with the presence of forskolin but in the absence of ELX-02.

All subsequent statistical analyses were performed using SPSS software (version 30.0.0, IBM). Group comparisons were conducted using Kruskal–Wallis tests, followed by Bonferroni-corrected Mann–Whitney U tests for pairwise comparisons, as most variables were not normally distributed. Paired responses across different ELX-02 concentrations were compared using the Wilcoxon signed-rank test. Correlations were assessed using the Spearman correlation coefficient. Enrichment of individual PTC variants among top-responding PDIOs was evaluated using 2×2 contingency tables, with odds ratios (ORs) and 95% confidence intervals calculated; significance was determined by Pearson Chi-square or Fisher’s Exact Test for small expected counts (cells <5), based on observed counts. All tests were two-sided, with statistical significance set at 0.05. No prior power analysis was performed. Graphs were made using GraphPad Prism (version 10.6.1).

## Results

### Study and subject overview

In the primary screen, PDIOs from 206 subjects carrying at least one nonsense variant were included. A full overview of the study design is provided in **Figure 1A**. Across this cohort, 48 distinct nonsense variants were analysed. Their positions within the CFTR mRNA are shown in **Figure 1B**, underlining the inclusion of nonsense variants throughout the whole CFTR gene.

**Figure 1.**
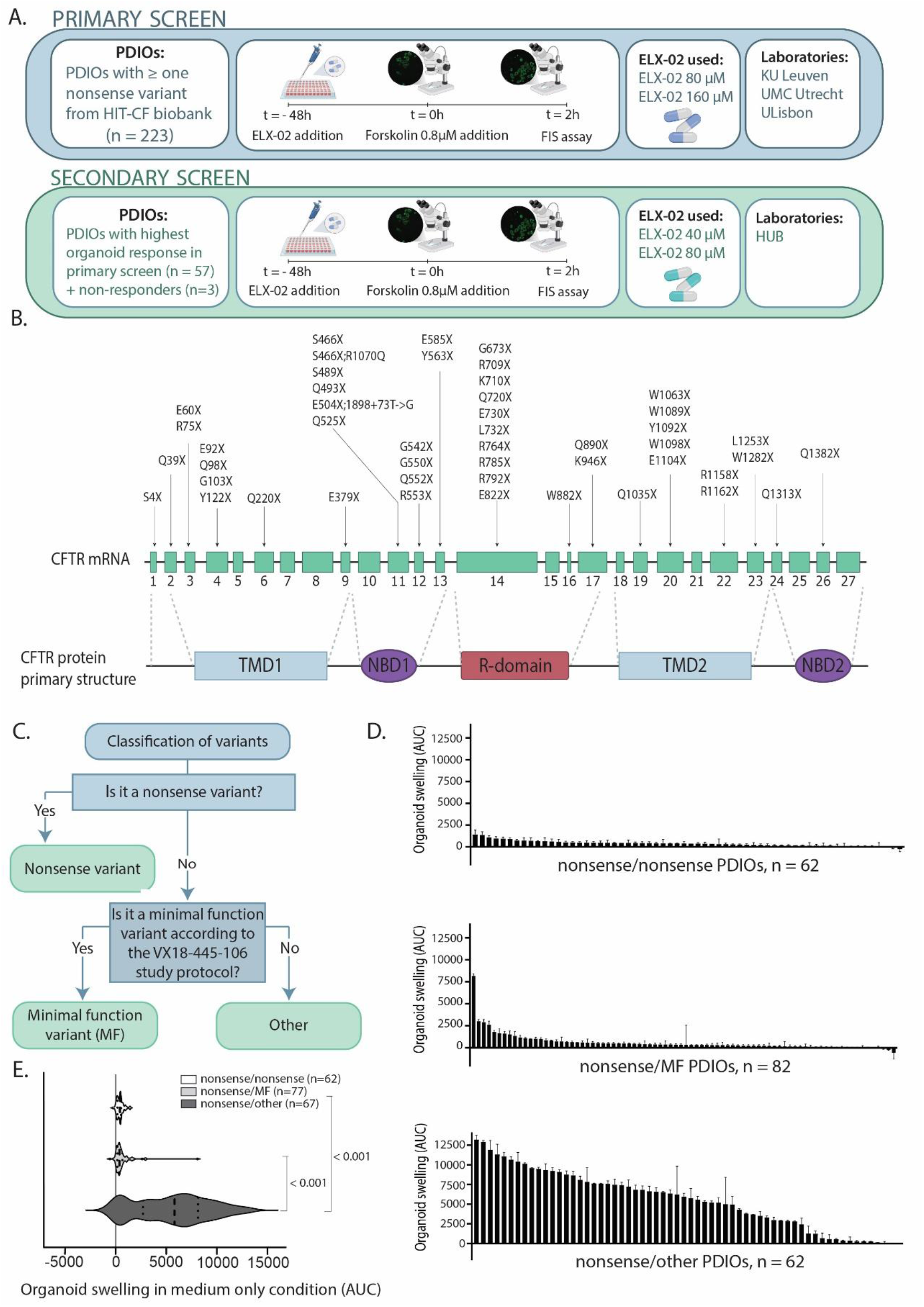
ELX-02 screening study design, CFTR nonsense variant distribution, genotype classification, and residual TR function. Schematic overview of the study design, including the number of included patients, the drug concentrations tested, the corresponding FIS assays performed, and the laboratories that conducted the screens (1A). Visualisation of the CFTR gene with the position of included nonsense variants, along with the corresponding CFTR primary protein structure (1B). Classification of genotypes into predefined categories used for all subsequent analyses. Variants were categorized as minimalction (MF) variants when listed as such in the VX18-445-106 study protocol (Study-VX18-445-106-Eligible-Mutationsne-2019.pdf) (1C). Waterfall plot of residual CFTR function measured by the FIS assay (2-hour swelling with medium only) subjects according to genotype categories (1D). Violin plot of residual function across the three genotype groups (1E).Biorender was used in the generation of Figure 1A

All variants reported in this study were classified into three categories: 1) nonsense variants, 2) minimal function (MF) variants, 3) other variants. Classification proceeded as follows: first all the nonsense variants according to the CFTR1 database nomenclature were assigned to the nonsense category (http://www.genet.sickkids.on.ca/). Next, all remaining (non-nonsense) variants were categorized according to the VX-445/TEZ/IVA eligibility list: variants designated as minimal-function in this document were classified as MF (24), and all others were categorized as “other”. An overview of this classification is provided in **Figure 1C**. Based on these criteria, PDIOs were grouped into three genotype categories: nonsense/nonsense (n = 62), nonsense/MF (n = 82), and nonsense/other (n = 62). Residual CFTR function was assessed by FIS over 2 hours in the absence of ELX-02 as shown for each PDIO in **Figure 1D**. Within the nonsense/MF group, one PDIO with genotype G542X/R1066C displayed unexpectedly high residual function, representing a notable outlier.

Residual CFTR function differed significantly across the three genotype groups (H = 74.06, df = 2, p < 0.001). No difference was observed between the nonsense/nonsense and nonsense/MF groups, whereas the nonsense/other group exhibited higher residual function than both the nonsense/nonsense and nonsense/MF groups, as shown in **Figure 1E**.

### Genotype-dependent response to ELX-02 in PDIOs primary screen

We next set out to compare genotype-dependent responses to ELX-02. **Figure 2A** and **2B** show the residual function-corrected 0.8 µM FIS response to ELX-02 at 80 µM in PDIOs belonging to the nonsense/nonsense and nonsense/MF genotype groups, respectively. Only PDIOs with a response above 1384 AUC are shown, corresponding to the benchmark of the mean response to ELX-02 at 80 µM in G542X-homozygous PDIOs. Responses below this benchmark are considered not clinically relevant and are not shown in the figure. The residual function-corrected 0.8 µM FIS response to ELX-02 at 80 µM in PDIOs belonging to the nonsense/other genotype group is shown in **Figure S2**.

**Figure 2.**
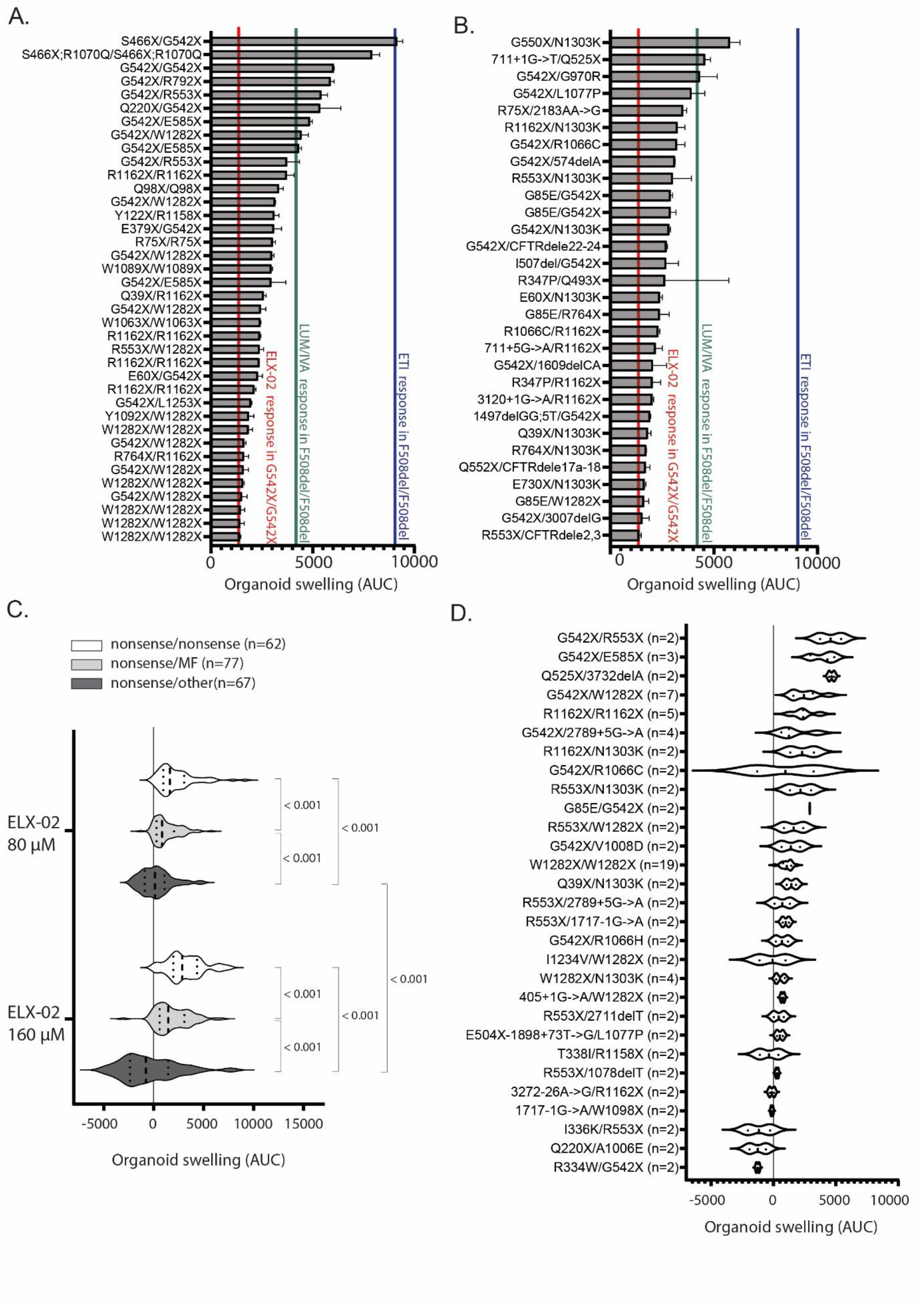
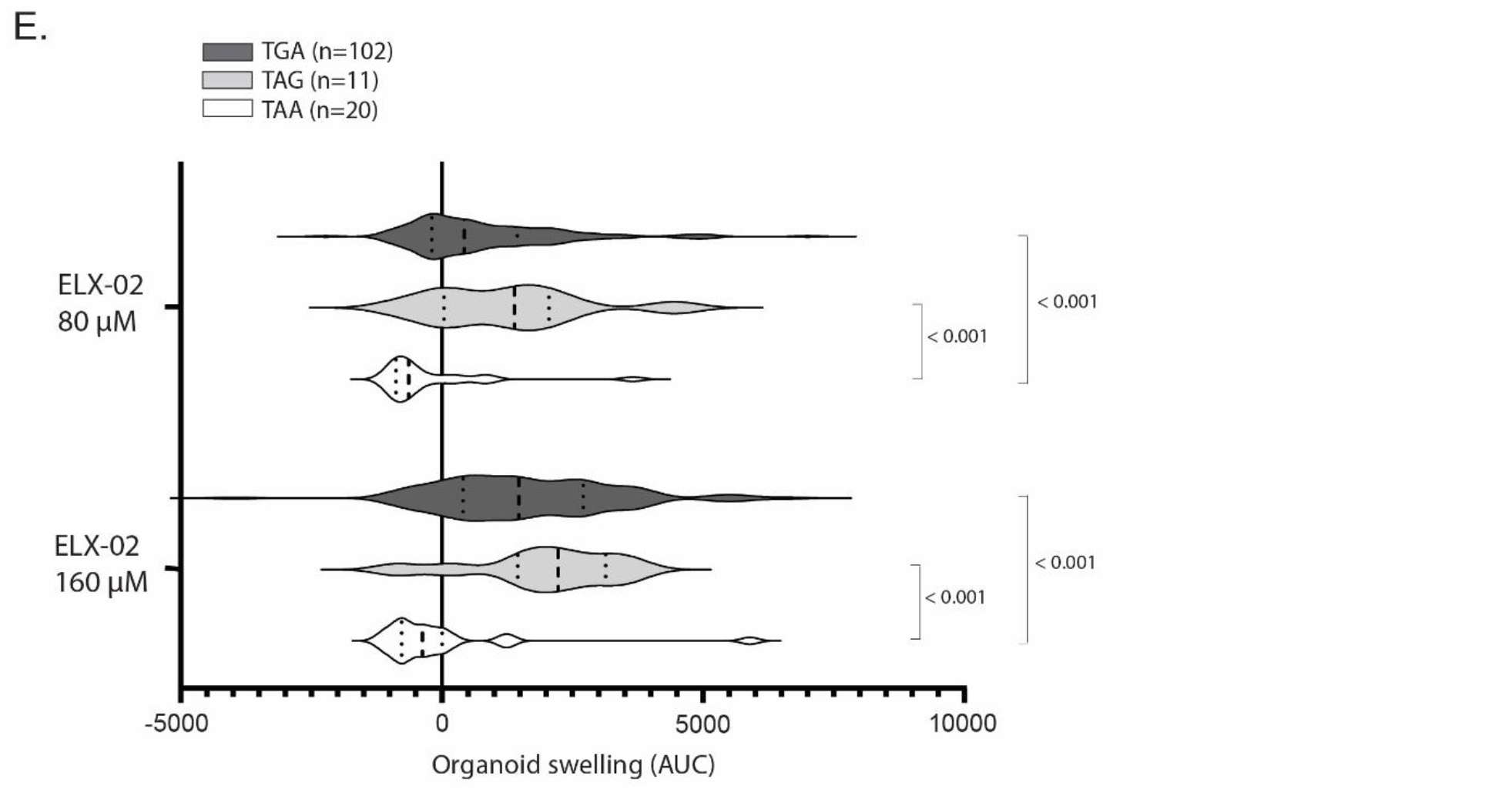
Genotype-dependent response to ELX-02 in PDIOs primary screen. Waterfall plot for residual function-corrected µM forskolin FIS responses to ELX-02 at 80 µM for PDIOs harboring two nonsense variants (2A) and for PDIOs harboring e nonsense variant and one minimal function variant (2B). Three benchmarks are included: 1. Mean ELX-02 80 µM residual ction-corrected FIS response of G542X/G542X PDIO reference line at 0.8 µM forskolin indicated in red (1384 AUC); 2. an lumacaftor/ivacaftor (LUM/IVA) residual function-corrected FIS response of F508del/F508del PDIO reference line at µM forskolin indicated in green (4187 AUC); 3. Mean elexacaftor/tezacaftor/ivacaftor (ETI) residual function-corrected S response of F508del/F508del PDIO reference line at 0.8 µM forskolin indicated in green (9052 AUC). Error bars represent M (n = 2). Only PDIOS with ELX-02 corrected responses above the mean G542X/G542X ELX-02 at 80 µM response 84 AUC) were included in both 2A and 2B graph. Responses below this benchmark are considered not clinically relevant d are not shown in the figure. Violin plot of the three categories across the two conditions used in the primary screen (ELX-160 µM residual-function corrected, ELX-02 80 µM residual function-corrected) (2C). Violin plot of all genotypes with re than one PDIOs in the screen (mean AUC FIS ELX-02 80 µM) (2D). Violin plot of residual function-corrected swelling uced by ELX-02 at 80 and 160 µM according to type of premature termination codon (PTC) in PDIOs with one nonsense d one minimal function variant or two nonsense variants with the same type of PTC (2E).

Within the nonsense/nonsense group (n = 62), one PDIO exhibited a response exceeding the mean response to elexacaftor/tezacaftor/ivacaftor (ETI) in F508del-homozygous PDIOs, and eight PDIOs showed responses exceeding the mean LUM/IVA response benchmark. Thirty-eight PDIOs exceeded the mean response to ELX-02 in G542X-homozygous PDIOs as shown in **Figure 2A**. In the nonsense/MF group (n = 77), three PDIOs demonstrated mean responses above the mean LUM/IVA benchmark and thirty PDIOs showed responses exceeding mean ELX-02 response in G542X-homozygous PDIOs, as shown in **Figure 2B**.

Inspection of individual responses revealed that the two highest-responding PDIOs both carried the S466X variant, with the S466X/G542X genotype showing a response comparable to the mean ETI response observed in F508del-homozygous PDIOs.

Among the top 10% responders, three nonsense variants, S466X, G542X and Q525X, showed enrichment (S466X: OR 1.11, 95% CI 0.96 – 1.27, p = 0.010; G542X: OR 4.54, 95% CI 1.79–11.51, p < 0.001; Q525X: OR 1.17, 95% CI 0.98 – 1.39, p < 0.001), whereas other nonsense variants, including G550X, R553X and E585X, were also observed among the top 10% responders but were not enriched. Interestingly, all these nonsense variants were located within the NBD1 domain. In contrast, PDIOs harboring W1282X were underrepresented among the top 10% responders (OR 0.14, 95% CI 0.018 – 1.06, p = 0.030) indicating that this variant is associated with lower ELX-02 responsiveness.

Swelling upon ELX-02 at 80 µM differed significantly across the three genotype categories (**Figure 2C**). The residual function-corrected FIS responses to ELX-02 at 80 µM were highest in PDIOs carrying two nonsense variants, compared with PDIOs carrying one nonsense and one MF variant. Both the nonsense/nonsense and nonsense/MF groups showed significantly higher residual function-corrected FIS responses than the nonsense/other group (**Figure 2C**). Using ELX-02 at 160 µM, similar responses in both the nonsense/nonsense and nonsense/MF were found compared to the response to ELX-02 at 80 µM. However, in PDIOs with nonsense/other genotype, ELX-02 at 160 µM concentration reduced FIS responses compared to 80 µM (**Figure 2C**). Negative FIS values indicate that FIS after exposure to ELX-02 was lower than residual function. At 160 µM ELX-02, more negative responses were observed. Organoid morphology in PDIOs with high residual function and subsequent reduced swelling upon adding ELX-02 is shown in **Figure S1**.

Although some of the PDIOs harbor identical genotypes, the majority carry unique compound-heterozygous genotypes. Therefore, our study has a theratyping character rather than a genotyping approach. In **Figure 2D**, the violin plots of genotypes with at least two available PDIOs reveal higher variability for G542X/R1066C and I1234V/W1282X, whereas genotypes such as Q525X/3732delA, G85E/G542X and 1717-1G>A/W1088X show minimal variability. ELX-02 responses at 80□µM did not differ between PDIOs with nonsenses close to the downstream exon–intron junction (≤55□nt) and those further away (>55□nt), suggesting that downstream junction distance alone does not predict readthrough efficacy (Mann-Whitney U = 3545, p = 0.075).

Finally, responses were analyzed according to the type of nonsense codon in PDIOs carrying either two nonsense variants with the same nonsense codon or one nonsense variant and one MF variant. Most nonsense variants introduced a TGA stop codon (n = 102), compared with TAA (n = 20) and TAG (n = 11). While residual function did not differ between nonsense categories, ELX-02-induced responses at both 80 µM and 160 µM were significantly lower in PDIOs harboring TAA nonsense codons than in those with TAG and TGA (**Figure 2E**).

### Validation of genotype-dependent differential responses to ELX-02

In the secondary screen, 57 PDIOs with the highest response and 3 PDIOs with no response as control were selected, including 31 nonsense/nonsense, 21 nonsense/MF and 8 nonsense/other PDIOs, and re-screened at a different institute (HUB). ELX-02 was tested at lower concentrations (40 µM and 80 µM) than in the primary screen, reflecting clinically relevant ELX-02 plasma levels (6, 25).

Residual function and residual function-corrected FIS responses to ELX-02 at 80 µM and at 40 µM did not differ significantly across the three genotype groups (**Figure S3**). **Figures 3A** and **3B** show the individual responses to ELX-02 at 80 µM and at 40 µM for nonsense/nonsense and nonsense/MF, respectively. Residual function-corrected FIS responses to ELX-02 at 80 µM were significantly higher than at 40 µM (Wilcoxon signed-rank test, p < 0.001). **Figure 3C** illustrates intragenotype variation for genotypes with two or more PDIOs. Responses were relatively consistent for R1162X/N1303K, W1282X/W1282X, and G542X/E585X, whereas responses in G542X/R553X were more heterogeneous, indicating higher intragenotype variability in this case.

**Figure 3.**
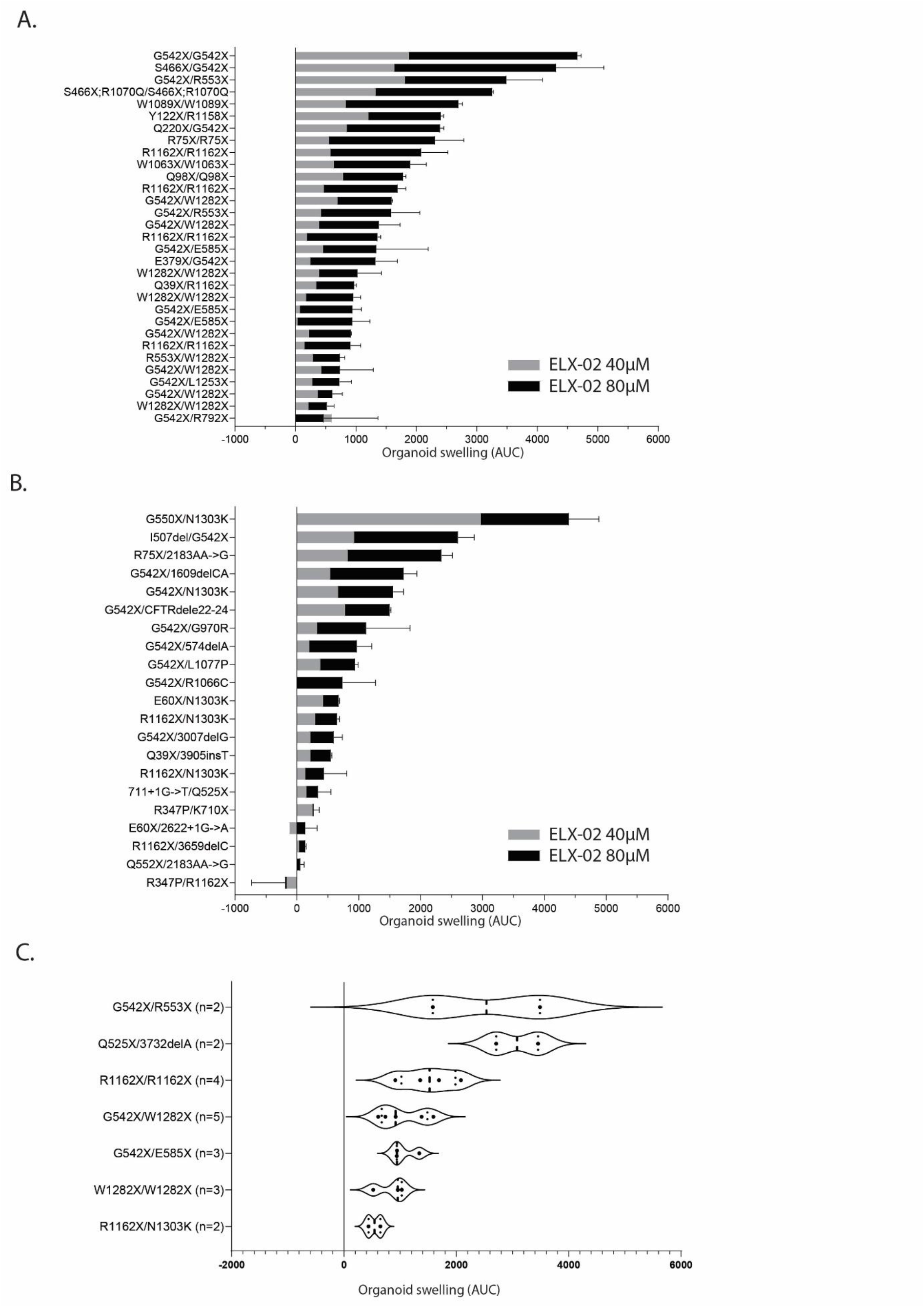
Genotype-dependent response to ELX-02 in PDIOs secondary screen. Waterfall plot for residual function-corrected 0.8 µM forskolin FIS responses to ELX-02 at 40 µM and 80 µM for PDIOs harboring two nonsense variants (3A) and for PDIOs harboring one nonsense variant and one minimal function variant (3B). Violin plot of all genotypes with more n one PDIOs in the screen (mean AUC FIS ELX-02 80 µM) (3C).

Overall individual response patterns were comparable between primary and secondary screen, such as S466X-containing genotypes consistently ranking among highest responders whereas PDIOs carrying W1282X variants again clustered among the lower responders. A comparison between results from the primary screen to the secondary screen are shown in **Figure S4**.

## Discussion

We describe FIS responses to the readthrough agent ELX-02 in a large cohort of PDIOs from people with CF harboring a broad range of nonsense variants, extending prior PDIO studies focused mostly on G542X, for which ELX-02 failed to demonstrate clinical benefit (6, 8, 13). Responses to ELX-02 varied markedly between subjects, ranging from effects that approached those observed in F508del/F508del PDIOs treated with LUM/IVA, to minimal responses. Overall, maximal FIS responses were modest and remained at the lower end of the range observed for approved CFTR modulators, consistent with the limited clinical efficacy reported for ELX-02 to date (6, 13). These findings indicate that ELX-02 monotherapy is likely insufficient for robust functional rescue, underscoring the need to better understand the determinants of readthrough efficacy and to develop more effective strategies.

PDIO responses to ELX-02 were dose-dependent and higher in PDIOs carrying two nonsense variants compared with those carrying a single nonsense variant in combination with a residual or minimal function allele, as expected based on target availability. Among individual genotypes, G542X PDIOs consistently showed higher responses than most other nonsense variants. Elevated responses were also observed for S466X, G550X and R553X, which are notably all located within NBD1. However, responsiveness was not dependent on the distance of the nonsense variant to the nearest intron or the canonical ±55 base pair region associated with nonsense-mediated decay (26), although it should be noted that we did not directly quantify mutant transcript abundance (e.g., by qPCR), and therefore cannot fully exclude a contribution of NMD-mediated reduction of mutant CFTR mRNA to the variability in ELX-02 responsiveness observed across variants. One possible explanation for the higher responsiveness observed among several NBD1-located variants is that amino acid substitutions generated during readthrough are better tolerated at specific NBD1 residues. Notably, variants such as S466X are located outside the canonical ATP-binding and catalytic motifs of NBD1, suggesting that successful readthrough may generate full-length CFTR proteins that retain substantial function despite incorporation of a non-native amino acid. Consequently, nonsense variants at these positions may be less sensitive to the identity of the incorporated amino acid, although local sequence context surrounding the PTC may also contribute, as has been suggested in the context of tRNA-based suppression therapies and may similarly apply to readthrough approaches (27). More likely, however, responsiveness is determined by the functional properties of the resulting readthrough-derived missense CFTR protein. This concept is exemplified by G550X, for which the putative readthrough product G550W has been shown to generate a highly functional CFTR protein (28). Stratification by stop codon identity revealed higher median responses for TGA codons, the most frequent stop codon in CFTR, and lower responses for TAA codons. This is consistent with previous findings demonstrating that translation termination fidelity and readthrough efficiency depend, among other factors, on stop codon identity, with termination efficiency ranked as TAA > TAG > TGA and corresponding readthrough efficiency as TGA > TAG > TAA (29). Nonetheless, this association was not absolute, as exemplified by the TAA variant Q525X, which was among the strongest responders. Thus, neither stop codon identity nor nonsense position alone appears sufficient to predict readthrough efficacy. Instead, responsiveness likely reflects the combined effects of transcript availability, local sequence context, readthrough efficiency, and the functional consequences of the amino acid incorporated during readthrough (30). Future studies combining quantitative proteomics allowing identification of incorporated amino acids, and functional assessment of corresponding missense variants may help disentangle the relative contribution of these mechanisms.

Despite the identification of responsive genotypes, the overall magnitude of rescue remained limited. To interpret these responses in a clinical context, three benchmarks can be considered. The lowest benchmark is the mean ELX-02-induced FIS response in G542X/G542X PDIOs, since clinical trials with ELX-02 failed to demonstrate clinically meaningful improvements in lung function or biomarkers, either as monotherapy or in combination with a potentiator (6, 13). PDIOs response falling below this threshold are therefore unlikely to confer any clinical benefit. The intermediate benchmark is the LUM/IVA-induced FIS response in F508del/F508del PDIOs, a modulator that demonstrated only modest improvements in FEV[ of approximately three percent predicted in clinical trials, detectable only in large patient cohorts (31). PDIOs with responses between the ELX-02 and LUM/IVA benchmarks thus represent a zone of uncertain clinical relevance. The highest benchmark is the ETI-induced FIS response in F508del/F508del PDIOs, which corresponds to robust clinical benefit. PDIOs exceeding this threshold are therefore most likely to show a meaningful clinical response. The relatively low range of ELX-02–mediated rescue in PDIOs therefore provides a plausible explanation for the absence of clear clinical benefit observed thus far. Although ELX-02 was assessed as a standalone readthrough agent to enable direct comparison of variant-specific efficacy, future studies could evaluate the combination of ELX-02 or follow-up readthrough molecules with CFTR modulators, as these may further rescue the function of readthrough-generated CFTR protein increasing therapeutic benefit.

Variability within genotypes was observed, consistent with previous work, possibly reflecting the heterogeneity in clinical responses among people with CF carrying identical CFTR variants treated with CFTR modulators, likely due to the influence of genetic modifiers and other patient-specific factors (8, 32).

This work underlines the need for further improvements on small molecule-based readthrough strategies. Although combinatorial approaches could theoretically enhance rescue through corrector or potentiator mechanisms, prior work, including our own, indicates that ELX-02 combined with ETI does not achieve sufficient functional rescue to expect clinical benefit (20). Future studies investigating whether the efficacy of ELX-02 can be further strengthened by newer and more effective modulators will be of interest, as will continued progress in the chemical optimization of readthrough compounds and the development of nonsense-mediated decay inhibitors.

Despite limited ELX-02 efficacy, these findings underscore the need for informed patient selection when designing clinical trials for readthrough therapies, given the large heterogeneity in responses to readthrough compounds. PDIO-based functional screening suggests that not all nonsense variants respond equally, and that responses near the detection threshold can vary substantially. Ignoring this heterogeneity may dilute treatment effects and increase the risk of negative trial outcomes, even when a subset of patients could derive meaningful benefit in future clinical trials. Incorporating PDIO-based preclinical readouts into trial design enables rational prioritization of patients with more responsive nonsense variants, increasing the likelihood of detecting true clinical efficacy and supporting precise, mechanism-informed trial stratification, as recently demonstrated in the context of the CHOICES trial (ClinicalTrials.gov NCT064685270; EudraCT 2022-500410-26-01). In this context, systematic evaluation of novel therapeutic strategies across individual nonsense variants in a relevant preclinical model such as PDIOs is therefore essential.

## Acknowledgements

This is a study connected to the HIT-CF Organoid Study group, which consists of the following people:

Helmut Ellemunter, Katharina Niedermayr (Medical University of Innsbruck, Cystic Fibrosis Center, Department of Child and Adolescent Health, Univ. Clinic for Paediatrics III, Innsbruck, Austria), Anabela S. Ramalho, Francois Vermeulen (Katholieke Universiteit Leuven, Leuven, Belgium), Elke De Wachter, Eef Vanderhelst (CF Center, Universitair Ziekenhuis Brussel, Vrije Universiteit Brussel, Brussels, Belgium), Tereza Dousova, Lucie Borek-Dohalska (Department of Pediatrics, University Hospital Motol and Second Faculty of Medicine, Charles University, Prague, Czechia), Marianne Skov, Tacjana Pressler (CF Center Copenhagen, Rigshospitalet, Copenhagen University Hospital, Copenhagen, Denmark), Edward F Nash, Marcus Mottershead (University Hospitals Birmingham NHS Foundation Trust, Birmingham, UK), Daniel Peckham (Leeds Teaching Hospitals NHS Trust, Leeds, UK), Mary Carroll (Adult Cystic Fibrosis Unit, University Hospital Southampton NHS Foundation Trust, Southampton, UK), Janisha Patel (Department of Hepatology, University Hospital Southampton NHS Foundation Trust, Southampton, UK), Helen Barker, Marlene Taveira (Royal Papworth Hospital NHS Foundation Trust, Cambridge, UK), Nicholas J. Simmonds (Adult Cystic Fibrosis Center, Royal Brompton Hospital, London, UK; National Heart and Lung Institute, Imperial College London, UK), Kinesh Patel (Imperial College London, UK; Royal Brompton Hospital, London, UK), Helen L Barr, Jessica Longmate (Wolfson Cystic Fibrosis Center, Department of Respiratory Medicine, Nottingham University Hospitals NHS Trust, Nottingham, UK), Michael D Waller (King’s College Hospital NHS Foundation Trust, Adult Cystic Fibrosis and Respiratory Medicine, London, UK), Peter J. Barry (Manchester University NHS Foundation Trust, Manchester Adult Cystic Fibrosis Center, Wythenshawe Hospital, Manchester, UK; Division of Infection, Immunity and Respiratory Medicine, School of Biological Sciences, University of Manchester, Manchester, UK), Dipesh H Vasant (Gastroenterology, Wythenshawe Hospital, Manchester University NHS Foundation Trust, Manchester, UK; Division of Diabetes, Endocrinology and Gastroenterology, University of Manchester, Manchester, UK), Michael Fayon, Julie Macey (University Hospital, Hôpital Pellegrin-Enfants, Pediatric Cystic Fibrosis Reference Center (CRCM), Center d’Investigation Clinique (CIC 1401), Bordeaux, France), Nadine Desmazes-Dufeu (Bordeaux University, Center de Recherche Cardio-Thoracique de Bordeaux, Bordeaux, France), Marlène Murris, Marion Dupuis (Service de Pneumologie-Allergologie CRC Mucoviscidose Adulte, Center de Transplantations Pulmonaire, Hôpital Larrey, Toulouse, France), Raphaël Chiron, Alexandre Coudrat (Cystic Fibrosis Center, Hôpital Arnaud de Villeneuve, Center Hospitalier Universitaire de Montpellier, Univ Montpellier, Montpellier, France), Isabelle Durieu, Quitterie Reynaud (Adult Cystic Fibrosis Care Center, Hospices Civils de Lyon, EA HESPER 7425, Université de Lyon, Lyon, France), Silke van Koningsbruggen, Jan-Christoph Thomassen (CF Center and Experimental Pulmonology, Children’s Hospital, Faculty of Medicine, University of Cologne, Cologne, Germany), Susanne Naehrig (Medizinische Klinik V, Cystic Fibrosis Center for Adults, LMU University of Munich, Munich, Germany), Felix C. Ringshausen, Annette Sauer-Heilborn (Department of Respiratory Medicine, Hannover Medical School (MHH) and Biomedical Research in End-stage and Obstructive Lung Disease Hannover (BREATH), German Center for Lung Research (DZL), Hannover, Germany), Simon Y. Graeber, Marcus A. Mall (Department of Pediatric Pulmonology, Immunology and Critical Care Medicine and Cystic Fibrosis Center, Charité-Universitätsmedizin Berlin; Berlin Institute of Health (BIH); German Center for Lung Research (DZL), Berlin, Germany), Sivagurunathan Sutharsan, Matthias Welsner (Division of Cystic Fibrosis, Department of Pulmonary Medicine, University Medicine Essen – Ruhrlandklinik, Essen, Germany), Michael Lorenz (Cystic Fibrosis Center /Pediatric Pneumology, University of Jena, Jena, Germany), Sabine Wege (Department of Pulmonology and Critical Care Medicine, Thoraxklinik at the University of Heidelberg, Heidelberg, Germany), Michael Wilschanski, Malena Cohen-Cymberknoh (Center for Cystic Fibrosis, Hadassah Hebrew University Medical Center, Jerusalem, Israel), Dario Prais, Meir Mei-Zahav (Pulmonary Institute, Schneider Children’s Medical Center of Israel; Sackler Faculty of Medicine, Tel Aviv University, Israel), Vincenzina Lucidi, Fabiana Ciciriello (Pediatric Hospital Bambino Gesù, Rome, Italy), Vito Terlizzi, Giovanni Taccetti (Cystic Fibrosis Regional Center of Tuscany, Anna Meyer Children’s University Hospital, Firenze, Italy), Cresta Federico, Carlo Castellani (Cystic Fibrosis Center, IRCCS Istituto Giannina Gaslini, Genoa, Italy), Giovanna Pizzamiglio, Leonardo Terranova (Fondazione IRCCS Ca’ Granda Ospedale Maggiore Policlinico, Internal Medicine Department, Respiratory Unit and Adult Cystic Fibrosis Center, Milan, Italy), Carla Colombo, Arianna Bisogno (Cystic Fibrosis Center, Fondazione IRCCS Ca’ Granda, Ospedale Maggiore Policlinico, Milan, Italy), Paola Melotti, Davide Treggiari (Azienda Ospedaliera Universitaria Integrata Verona, Verona, Italy), Claudio Sorio (University Dept. of Medicine, General Pathology Division, Univ. of Verona, Verona, Italy), Barbara Messore, Sara Demichelis (Azienda Ospedaliero Universitaria San Luigi Gonzaga, Adult Cystic Fibrosis Center, Pulmonology Div., Turin, Italy), Dorota Sands, Lukasz Wozniacki (Institute of Mother and Child, Cystic Fibrosis Department, Warsaw, Poland), Celeste Barreto, Carolina Gonçalves (Departamento de Pediatria, Hospital de Santa Maria (CHULN), Centro Académico de Medicina de Lisboa, Lisboa, Portugal), Margarida Amaral, Iris Silva (BioISI – Biosystems & Integrative Sciences Institute, University of Lisboa, Lisbon, Portugal), Silvia Gartner (Pediatric Pulmonology and Cystic Fibrosis Unit, Hospital Universitari Vall d’Hebron, Barcelona, Spain), Antonio Alvarez Fernandez (Adults Cystic Fibrosis Unit, Hospital Universitari Vall d’Hebron, Vall d’Hebron Institut de Recerca (VHIR), Barcelona, Spain), Lena Hjelte, Isabelle de Monestrol (Stockholm CF Center, Karolinska University Hospital, Karolinska Institutet, Stockholm, Sweden), Marita Gilljam (Department of Respiratory Medicine, Sahlgrenska University Hospital, Gothenburg, Sweden), Per Hedenström (Unit of GI Endoscopy, Sahlgrenska University Hospital, Gothenburg, Sweden), Alexander Möller (University of Zurich, Switzerland), F. Singer (Division of Respiratory Medicine, Dept of Pediatrics, Inselspital, University of Bern, Bern, Switzerland), M. Bakker (Department of Pulmonology, Erasmus MC, University Medical Center, Rotterdam, The Netherlands), H.M. Janssens (Division of Respiratory Medicine and Allergology, Dept of Pediatrics, University Medical Center Rotterdam, Erasmus MC-Sophia, Rotterdam, The Netherlands), J. Altenburg (Department of Respiratory Medicine, Amsterdam University Medical Centres, University of Amsterdam, Amsterdam, The Netherlands), S.W.J. Terheggen-Lagro (Department of Pediatric Pulmonology, University Medical Center Utrecht, The Netherlands).

This work was supported by the European Union’s Horizon 2020 research and innovation program under grant agreement No. 755021.

