## Supplemental Table 1 & Supplemental Figures 1-3 for "A functional comparison of readthrough agent ELX-02 across a wide range of nonsense CFTR variants"

Supplementary material

VERTEX
THE SCIENCE of POSSIBILITY

### Eligible MF *CFTR* Mutations for a Phase 3 Study Evaluation the Pharmcokinetics, Safety, and Tolerability of VX-445/TEZ/IVA Triple Combination Therapy in Cystic Fibrosis Subjects 6 Through 11 Years of Age

The below list includes currently eligible minimal function mutations for the VX18-445-106 study as of June 2019 (protocol version 2.0):

#### Non-exhaustive List of Minimal Function *CFTR* Mutations Eligible for VX18-445-106

| | | | | |
|---|---|---|---|---|
| Q2X | L218X | Q525X | R792X | E1104X |
| S4X | Q220X | G542X | E822X | W1145X |
| W19X | Y275X | G550X | W882X | R1158X |
| G27X | C276X | Q552X | W846X | R1162X |
| Q39X | Q290X | R553X | Y849X | S1196X |
| W57X | G330X | E585X | R851X | W1204X |
| E60X | W401X | G673X | Q890X | L1254X |
| R75X | Q414X | Q685X | S912X | S1255X |
| L88X | S434X | R709X | Y913X | W1282X |
| E92X | S466X | K710X | Q1042X | Q1313X |
| Q98X | S489X | Q715X | W1089X | Q1330X |
| Y122X | Q493X | L732X | Y1092X | E1371X |
| E193X | W496X | R764X | W1098X | Q1382X |
| W216X | C524X | R785X | R1102X | Q1411X |

| | | | | |
|---|---|---|---|---|
| 185+1G>T | 711+5G>A | 1717-8G>A | 2622+1G>A | 3121-1G>A |
| 296+1G>A | 712-1G>T | 1717-1G>A | 2790-1G>C | 3500-2A>G |
| 296+1G>T | 1248+1G>A | 1811+1G>C | 3040G>C (G970R) | 3600+2insT |
| 405+1G>A | 1249-1G>A | 1811+1.6kbA>G | | 3850-1G>A |
| 405+3A>C | 1341+1G>A | 1811+1643G>T | 3120G>A | 4005+1G>A |
| 406-1G>A | 1525-2A>G | 1812-1G>A | 3120+1G>A | 4374+1G>T |
| 621+1G>T | 1525-1G>A | 1898+1G>A | 3121-2A>G | |
| 711+1G>T | | 1898+1G>C | | |

| | | | | |
|---|---|---|---|---|
| 182delT | 1119delA | 1782delA | 2732insA | 3791delC |
| 306insA | 1138insG | 1824delA | 2869insG | 3821delT |
| 365-366insT | 1154insTC | 1833delT | 2896insAG | 3876delA |
| 394delTT | 1161delC | 2043delG | 2942insT | 3878delG |
| 442delA | 1213delT | 2143delT | 2957delT | 3905insT |
| 444delA | 1259insA | 2183AA>G [a] | 3007delG | 4016insT |
| 457TAT>G | 1288insTA | 2184delA | 3028delA | 4021dupT |
| 541delC | 1343delG | 2184insA | 3171delC | 4022insT |
| 574delA | 1471delA | 2307insA | 3171insC | 4040delA |
| 663delT | 1497delGG | 2347delG | 3271delGG | 4279insA |
| 849delG | 1548delG | 2585delT | 3349insT | 4326delTC |
| 935delA | 1609del CA | 2594delGT | 3659delC | |
| 1078delT | 1677delTA | 2711delT | 3737delA | |

VERTEX
THE SCIENCE of POSSIBILITY

**Eligible MF *CFTR* Mutations for a Phase 3 Study Evaluation the Pharmcokinetics, Safety, and Tolerability of VX-445/TEZ/IVA Triple Combination Therapy in Cystic Fibrosis Subjects 6 Through 11 Years of Age**

The below list includes currently eligible minimal function mutations for the VX18-445-106 study as of June 2019 (protocol version 2.0):

**Non-exhaustive List of Minimal Function *CFTR* Mutations Eligible for VX18-445-106**

| | | |
|---|---|---|
| CFTRdele1 | CFTRdele16-17b | 991del5 |
| CFTRdele2 | CFTRdele17a,17b | 1461ins4 |
| CFTRdele2,3 | CFTRdele17a-18 | 1924del7 |
| CFTRdele2-4 | CFTRdele19 | 2055del9>A |
| CFTRdele3-10,14b-16 | CFTRdele19-21 | 2105-2117del13insAGAAA |
| CFTRdele4-7 | CFTRdele21 | 2372del8 |
| CFTRdele4-11 | CFTRdele22-24 | 2721del11 |
| CFTR50kbdel | CFTRdele22,23 | 2991del32 |
| CFTRdup6b-10 | 124del23bp | 3121-977_3499+248del2515 |
| CFTRdele11 | 306delTAGA | 3667ins4 |
| CFTRdele13,14a | 602del14 | 4010del4 |
| CFTRdele14b-17b | 852del22 | 4209TGTT>AA |

| | | | |
|---|---|---|---|
| A46D | V520F | Y569D | N1303K |
| G85E | A559T | L1065P | |
| R347P | R560T | R1066C | |
| L467P | R560S | L1077P | |
| I507del | A561E | M1101K | |

**Supplementary table 1. Minimal function CFTR variants used in this study.** In this study, CFTR variants were categorized as minimal function (MF) based on the list defined in the VX18-445-106 study protocol.

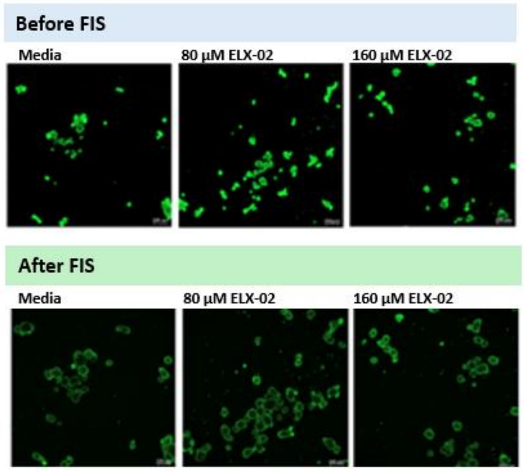


**Fig. S1. Organoid morphology in PDIOs with high residual function and inhibition of FIS by ELX-02.**

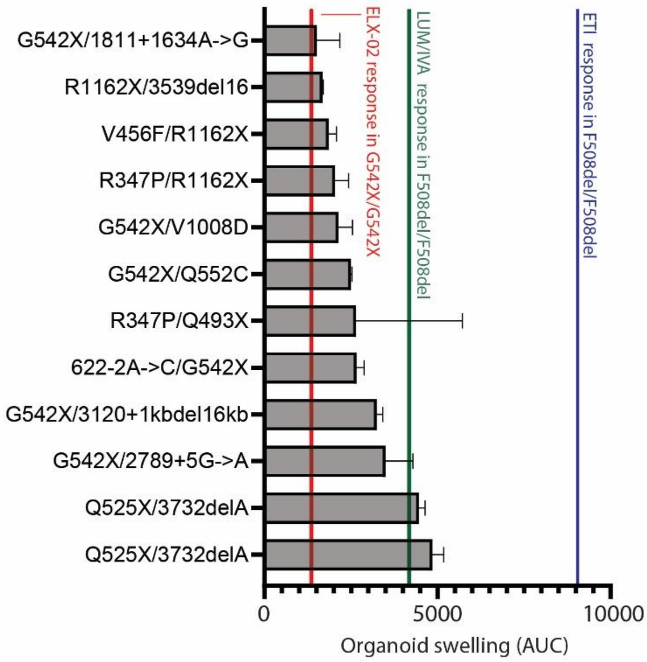


**Fig. S2. Waterfall plot for residual function-corrected 0.8µM forskolin FIS responses to ELX-02 at 80 µM for PDIOs harboring one nonsense variant and one "other" variant.** Three benchmarks are included: 1. Mean ELX-02 80 µM residual function-corrected FIS response of G542X/G542X PDIO reference line at 0.8 µM forskolin indicated in red (1384 AUC); 2. Mean lumacaftor/ivacaftor (LUM/IVA) residual function-corrected FIS response of F508del/F508del PDIO reference line at 0.8 µM forskolin indicated in green (4187 AUC); 3. Mean elexacaftor/tezacaftor/ivacaftor (ETI) residual function-corrected FIS response of F508del/F508del PDIO reference line at 0.8 µM forskolin indicated in green (9052 AUC). Error bars represent SEM (n = 2). Only PDIOs with ELX-02 corrected responses above the mean G542X/G542X ELX-02 at 80 µMresponse (1384 AUC) were included in the graph. Responses below this benchmark are considered not clinically relevant and are not shown in the figure.

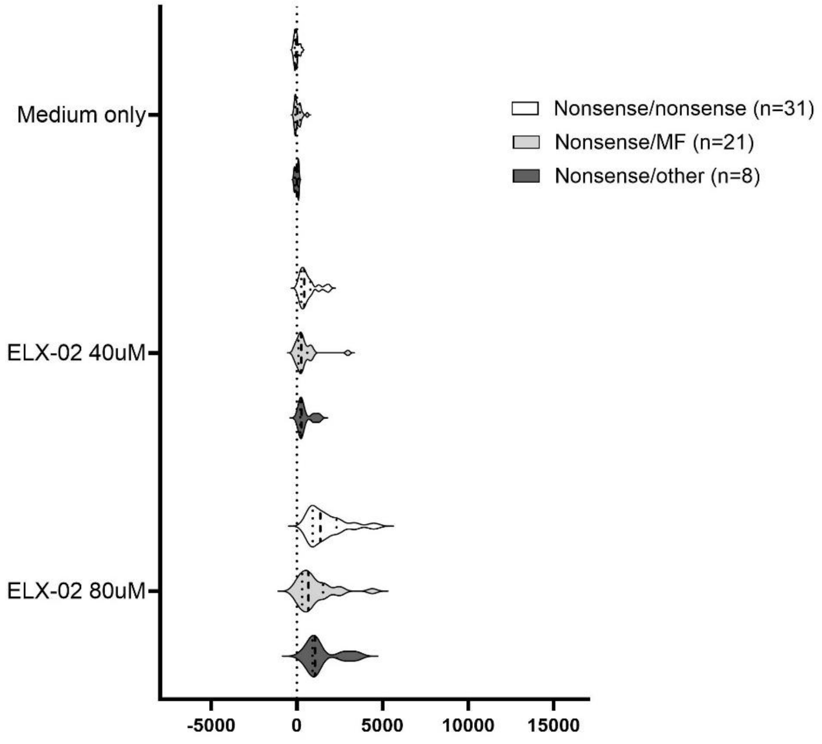


**Fig. S3. Genotype-dependent residual function and responses to ELX-02 in PDIOs in secondary screen (validation screen).** Violin plot of the three categories across the three conditions used in the secpndary screen (medium only, ELX-02 40 µM residual-function corrected, ELX-02 80 µM residual function-corrected).

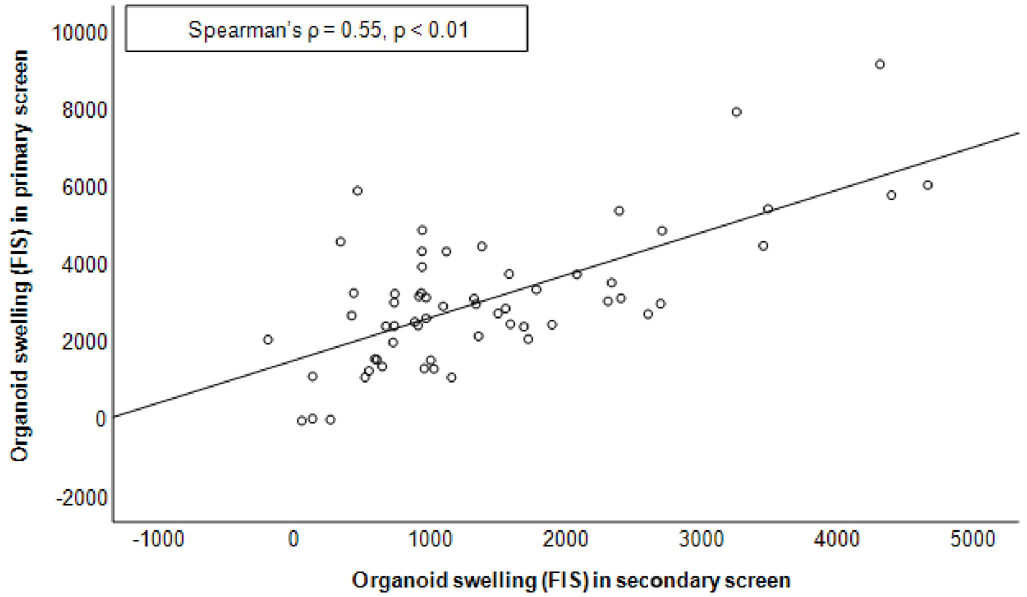


**Fig. S4. Scatterplot comparing FIS measurements between primary screen and secondary screen (validation screen).** FIS at 0.8 µM forskolin upon ELX-02 at 80 µM treatment. Each point represents an individual PDIO (n=60). Data are not normally distributed, therefore, the Spearman correlation coefficient ($\rho = 0.55$) and associated p-value ($p < 0.01$) are shown, indicating a moderate positive association between the primary and secondary screen responses.
